# Reorganization of molecular networks associated with DNA methylation and changes in the rearing environments of the house wren (*Troglodytes aedon*)

**DOI:** 10.1101/2021.05.04.442647

**Authors:** Bridgett M. vonHoldt, Rebecca Y. Kartzinel, Kees van Oers, Koen J.F. Verhoeven, Jenny Ouyang

## Abstract

Environmental change, such as increased rates of urbanization, can induce shifts in phenotypic plasticity with some individuals adapting to city life while others are displaced. A key trait that can facilitate adaptation is the degree at which animals respond to stress. This stress response has a heritable component and exhibits intra- and inter-individual variation. However, the mechanisms behind this variability and whether they might be responsible for adaptation to different environments are not known. Variation in DNA methylation can be a potential mechanism that mediates environmental effects on the stress response. We used an inter- and intra-environmental cross-foster experiment to analyze the contribution of DNA methylation to early-life phenotypic variation. We found that at hatching, urban house wren (*Troglodytes aedon*) offspring had increased methylation as compared to their rural counterparts, and observed plasticity in methylation as offspring aged, indicating developmental effects of the rearing environment on methylation. Differential methylation analyses showed that cellular respiration genes were differentially expressed at hatching and behavioral and metabolism genes were differentially expressed at fledgling. Lastly, hyper-methylation of a single gene (*CNTNAP2*) is associated with increased glucocorticoid levels. These differential methylation patterns linked to a specific physiological phenotype suggest that DNA methylation may be a mechanism by which individuals adapt to novel environments. Characterizing genetic and environmental influences on methylation is critical for understanding the role of epigenetic mechanisms in evolutionary adaptation.

## Introduction

With environments capable of rapid and dramatic change, phenotypic variation is increasingly investigated for molecules that contribute towards phenotypic plasticity through modified transcriptional control (e.g. Ledón-Rettig et al. 2012; Kilvitis et al. 2017; Jeremias et al. 2018; Viitaniemi et al. 2019; Taff et al. 2019). While plasticity occurs throughout an organism’s life (Pigliucci 2005), intergenerational plasticity also can occur when offspring phenotype results from parental environmental contexts (Chen et al. 2015; Verhulst et al. 2016; Venney et al. 2021). One main mechanism for transmission of intergenerational plasticity is through maternal effects on offspring development and phenotype (Groothuis & Schwabl 2008). The underlying genetic basis of variation in phenotypic traits consists of a combination of maternal or dominance effects (Lynch & Walsh 1998; Marshall & Uller 2007). Environmental effects may cause a single genotype to produce different phenotypes. When the genotype interacts with the environment (G × E), different genotypes may respond differently to this environmental variation, resulting in variable fitness consequences when genotypes are expressed in different environments. Therefore, disentangling the genetic basis, the environmental context, and the interaction between genetics and environmental variation is critical for understanding phenotypic variation and organismal evolution.

Epigenetic mechanisms can alter organism function without underlying changes in the DNA sequence, representing a possible mechanism for differences in the contribution of genetic and plastic variation to early-life traits (Bossdorf et al. 2008). The presence of 5-methylcytosine can immediately alter gene expression through a diverse array of mechanisms (e.g. splice sites, promoters, transcript stability), and this epigenetic methylation process commonly underlies environmentally-induced changes in gene expression (*e.g.*, Suzuki & Bird 2008; Goerlick et al. 2012; Ledón-Rettig et al. 2012; Heard et al. 2014; Pedersen et al. 2014; Lea et al. 2016; Sasagawa et al. 2017; Laubach et al. 2018). Early life experiences are one specific external cause that can change methylation within a lifetime with possible intra- and inter-generational stability (e.g. Heard et al. 2014; Lea et al. 2016; Rubenstein et al. 2016). Therefore, DNA methylation may be a potential mechanism for organismal acclimation and adaptation to novel environments.

Can a surveillance of methylation indicate the degree to which an individual’s early life environment predict future phenotype and local adaptation? To investigate this question, we conducted an inter-environment cross-fostering experiment using a free-living population of house wrens (*Troglodytes aedon*) in Nevada, U.S. with a subsequent surveillance of DNA methylation variation (Fig. 1A). Our design included a comparison between an urban site that has increased levels of noise, light and human density compounded by a highly fragmented landscape populated by non-native vegetation to a more natural, rural site (Baldan & Ouyang 2020). The anthropogenically altered landscape is characterized by decreased availability of natural high-quality food resources (Baldan & Ouyang 2020). Such areas are settled by younger, less-experienced individuals and consequently produce smaller-sized offspring than rural or natural habitat counterparts (Sprau et al. 2017; Sepp et al. 2017; Marzluff 2017). These characteristics reduce reproductive success for house wrens living in urbanized areas, as we have previously reported that their offspring are smaller at fledgling, with higher corticosterone levels ubiquitous across individuals (Ouyang et al. 2019). The rearing environment was identified as the causative agent as translocation of rural wrens to urban habitats increased corticosterone levels (Ouyang et al. 2019). Further, exposure of rural adult house wrens to experimental noise resulted in increased corticosterone levels, a trend that was not found in urban wrens, suggesting habituation (Davies et al. 2017).

**Figure 1.**
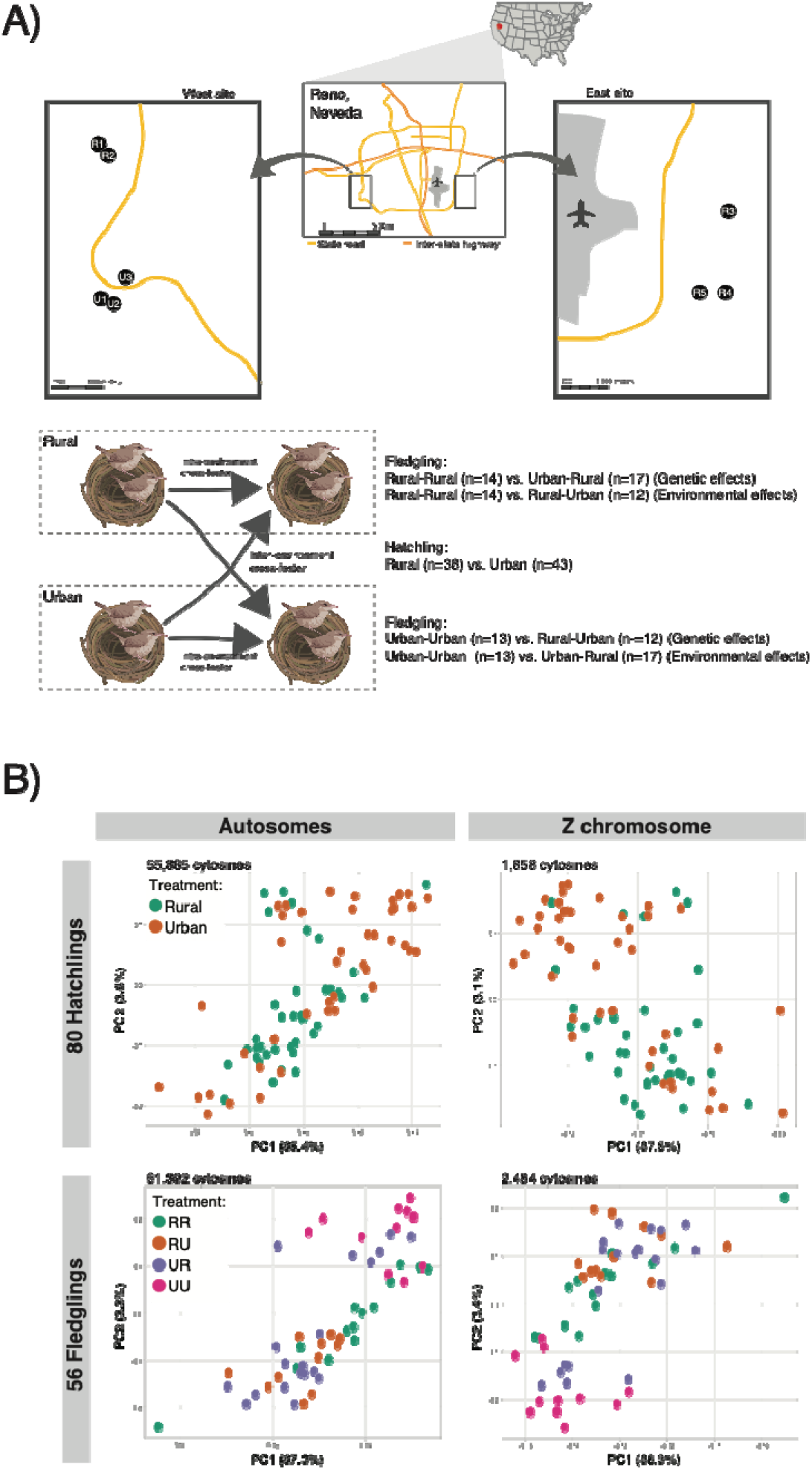
**A)** Map of geographic area from which source and recipient nests were selected in Rena, Nevada, USA. A schematic of the cross-fostering design depicts intra- and inter-environmental movement of individual wrens. Although eggs are depicted in this figure, hatchlings (on day 0) were translocated for cross-fostering. Sample sizes per cross-fostering treatment and age (hatchling, 0 days old; fledgling, 15 days old) after excluding samples that failed sequencing. Created with Biorender.com; **B)** Principal component analysis (PCA) of methylation data across CG motifs on the autosomes and Z chromosome derived from 80 house wrens sampled as hatchlings (day 0) and 56 fledglings (day 15) after cross-fostering manipulations. (Abbreviations: RR, rural hatchling that fledged from rural nest; RU, rural hatchling that fledged from urban nest; UR, urban hatchling that fledged from rural nest; UU, rural hatchling that fledged from urban nest)

Methylation in avian genomes follow the classic vertebrate paradigm of decreased transcription with increasing CpG methylation around transcription start sites (TSS) (Luo et al. 2011; Derks et al. 2016; Laine et al. 2016). With the aim to unravel the genetic and environmental determinants of DNA methylation variation, we had the following questions with respect to CpG methylation in house wrens from the cross-fostering experiment. First, does methylation differ with respect to the habitat location of the nest at an early life stage (hatching)? Second, is there a genetic signal on methylation at the time of fledging with respect to their hatching location? And finally, is there a signal of environmental stressors on methylation at the time of fledging with respect to their fledging location? We further combined DNA methylation sequencing data with a surveillance of glucocorticoid levels as a measurement of circulating levels of corticosterone hormones in blood. This multi-dimensional four-way crossfoster experiment allowed us to investigate the contributions of genetic and plasticity mechanisms that regulate offspring development.

## Materials and Methods

### Sample collection

We conducted our cross fostering study from May to July 2017 and 2018 at one urban location in Reno, Nevada, USA and one rural location in Sparks, Nevada, USA (for detailed description of the field sites, see Ouyang et al. 2019). (Fig. 1A). We monitored house wren nests regularly to determine the hatch date (day 0). On day 0 or 1, we obtained a blood sample from all chicks (2.1±0.02 mins; mean±S.E.). Chicks sampled on day 0 or 1 did not differ in their corticosterone levels (p>0.2). Any chick that was blood sampled past three minutes was not used in the study. After weighing all chicks, we individually marked them with toenail polish and toenail clipping for identification. Hatchlings were then translocated to a new nest to investigate the mechanistic and molecular basis of phenotypic variation in a 2×2 experimental design in a quartet (Fig. 1A). Internal controls for translation were obtained by fostering a hatchling originating from a rural nest with non-biological rural parents and a hatchling originating from an urban nest with non-biological urban parents (hereinafter referred to as RR and UU, respectively). We also translocated rural hatchlings to urban nests and urban hatchlings to rural nests (hereinafter referred to as RU and UR, respectively). During the crossfoster process, we transported the chicks in a heated nest box to either a rural or an urban nest that hatched on the same day (maximum transport time: 20mins, average transport time: 10mins). On day 8, we banded the nestlings with a uniquely numbered tarsal band. On day 15 (house wrens fledge around day 17 and reach asymptotic weight at day 15), we obtained another blood sample from all chicks (1.7 ± 0.02 mins) and weighed them to the nearest 0.1g.

### Corticosterone hormone collection

We used enzyme-linked immunosorbent assay kits (Enzo Life Sciences) following the manufacturer’s instructions. Please see Ouyang et al. (2019) for validation of this assay for house wrens. Based on optimization, we diluted plasma 1:40 in assay buffer with 0.5% steroid displacement reagent. We randomized samples across plates but an individual’s day 0 and day 15 plasma were always next to each other on the same plate. We included a standard curve on each plate that ranged from 32ng/mL to 20,000ng/mL. The assay sensitivity was 2.1pg/mL. To calculate intra- and inter-plate variation (CV), we included pooled house wren plasma, assayed in triplicate. The intra-plate CV was 9.8% and the inter-plate CV (5 plates) was 5.6%.

### Extraction of genomic DNA

We selected to conduct this temporal survey of blood-derived methylation variation as the individuals were not to be sacrificed and blood can be informative for several, but not all, tissue-specific patterns as well as for sites associated with expression (i.e. transcriptional start sites, TSS) (Husby 2020; Lindner et al. 2021). We extracted genomic DNA from avian whole blood, where both red (~99.5%) and white blood cells (0.5%) are nucleated (Scanes 2015), as per Qiagen DNeasy’s protocol with the following modifications (Qiagen). As we collected typically less than 20μL of blood, we added 180μL of 1x phosphate buffered saline (PBS) and 20μL of proteinase K to each sample. If the sample was represented by >20μL of whole blood, we added 360μL of 1x PBS directly into sample tube, mixed, and aliquoted equal volumes into separate 1.5mL tube, to which we added 20ul of proteinase K. The duplicate tubes were combined after the DNA extraction and spin column purification steps.

### Preparation, sequencing, and bioinformatics of bisulfite converted genomic libraries

We employed a reduced representation bisulfite sequencing (RRBS) approach using the *Msp1* restriction enzyme (Boyle et al. 2012) for whole-blood derived high molecular weight genomic DNA. Libraries were prepared following the NEBNext sample preparation kit (New England Biolabs). We added 1ng of enterobacteria phage lambda DNA to each genomic library as a non-methylated internal control for estimating bisulfite (BS) conversion downstream (e.g. Lea et al. 2015). We purified the genomic libraries and retained fragments between 100-400bp in size using AMPure beads, with subsequent treatment of the fragments with bisulfite to convert unmethylated cytosines with the low DNA input (1ng-2μg) protocol in the Qiagen EpiTect Fast Bisulfite Conversion kit (Qiagen). Converted DNA was subjected to 12 cycles of PCR amplification to enrich for adapter-ligated fragments with MyTaq Mix (Bioline Inc.), during which each sample was also barcoded with unique sequence tags. Libraries were randomly pooled with an average of 16 samples per lane for single-end (1×100nt) sequencing on an Illumina Novaseq 6000. We prepared the reference zebra finch genome (TaeGut2, GCA_000151805.2) with *bowtie2* in *BS-Seeker2* for BS-converted reads that were bounded from 50-300bp (Chen et al. 2010; Langmead 2010; Langmead & Salzberg 2012). We demultiplexed sequence pools using perfect sequence matches between expected and observed barcode sequence tags. FASTQ files were trimmed for low quality reads (Q<20) and adapter sequences clipped using *cutadapt 1.8.1* (Martin 2011), discarding reads that are <20bp in length. Trimmed reads were mapped to the reference with *bowtie2* and methylation called in *BS-Seeker2*. Cytosines were annotated as being either inter-genic or in a promoter (within 2Kb of transcriptional start site; Agirre et al. 2015), gene body (exon methylation; e.g. Bewick & Schmitz 2017), intron, or intergenic using gene annotations obtained from the Genes and Gene Predictions Tracks using the UCSC Table Browser for TaeGut2. Three TaeGut2 chromosomes (LG2, LG5, and LGE22) were unable to be annotated for genomic context due to a lack of information in the UCSC tracks. A cytosine may have membership in multiple categories; therefore, any cytosine that is minimally located within a single exon, it was considered as a gene body cytosine. Similarly, cytosines located within an intron but within 2Kb downstream of an exon of a second known gene were annotated as promoter-associated. Cytosines located both in an exon and inferred promoter were considered as a unique group. Inference of the regulatory consequences or *cis*-methylation is context dependent, where promoter methylation has been associated with transcriptional suppression while gene body methylation reduces transcriptional noise in other systems (Lister et al. 2009; Maunakea et al. 2010; Huh et al. 2013; Varley et al. 2013; Yang et al. 2014).

The methylation frequency (MF) per cytosine was calculated as the proportion of methylated cytosines out of the total number of methylated and unmethylated cytosine (sequenced as thymine) reads per site (Chen et al. 2010). We filtered to retain sites with a minimum of 10-fold sequencing coverage. We estimated the conversion efficiency by mapping each genomic library to the 48,502bp phage lambda linear genome (NC_001416.1) and assessed methylation levels of the converted lambda DNA. Conversion rates were estimated as [*1-average MF across the phage lambda genome*].

We further use the R v3.6.0 program *methylKit* (Akalin et al. 2012; R Core Team 2019) to apply a coverage filter to retain sites with a minimum of 10-fold and below a maximum read threshold of 95% percentile) that were sequenced in all individuals or samples identified to be analyzed. As cytosines found in the CG di-nucleotide motif are the predominant motif for vertebrate methylation (Suzuki & Bird 2008), we analyzed CG-motif cytosines only in all subsequent analyses. We also removed cytosines that were not variable across individuals. We assessed all samples together for cytosines present in all samples, as well as subsets of the samples for specific analyses (e.g. all rural hatchlings compared to all urban hatchlings, etc.). To assess the samples for quality and general patterns, we employed a principal component analysis (PCA) using *prcomp* function in R v3.6.0 (R Core Team 2019) to identify potential outliers for exclusion.

### Differential methylation analysis

We arranged samples into five different comparisons for identifying differentially methylated sites. (Fig. 1A) These comparisons were grouped into three sets for downstream analysis: 1) hatchlings in rural vs. urban environments (R vs. U); 2) fledglings that were hatched in the same environment but cross-fostered in different environments (RR vs. RU, UR vs. UU), to identify environmental influences in fledgling methylation patterns; and 3) fledglings that were hatched in different environments but fledged in the same environment (RR vs. UR, RU vs. UU), to identify the genetic influences in fledgling methylation patterns. We identified differentially methylated sites with the program *MACAU* that uses a binomial mixed model regression on methylation count data (Lea et al. 2015). PC coordinates were estimated using R’s *prcomp* function and the first two PCs were used as covariates in the linear mixed model to account for batch effects, along with the parental nest identity. We assumed full-sibling level of relatedness (r=0.5) for hatchlings in the same parental nest and no inter-nest relatedness (r=0). When no nest information is available, we assumed no relatedness.

To identify genomic sites and regions of differential methylation for further analysis, we used network analysis in the R package *igraph* (Csardi & Nepusz 2006). This analysis is based on the assumption that differential methylation that is most relevant will be 1) found in close proximity to other DMSs within the same treatment comparison (that is, a differentially methylated region or DMR); and/or 2) found in both treatment comparisons within a set (Fig. 1A). This holistic approach allows us to assess DMS lists from multiple comparisons simultaneously. We acknowledge that there is an inherent bias in physical clustering of cytosines given the reduced representation method of fragment selection. Our goal is to capture information at both the single-cytosine and ‘neighborhood’ level of methylation.

Within a set, DMS lists were pooled and treated as network vertices, with edges connecting DMSs within 40bp of each other. Clusters of DMSs (independent, maximal subgraphs with at least two vertices) were extracted from the networks for further analysis. We refer to these clusters, which are comprised of DMSs from one or more treatment comparisons, as *differentially methylated clusters*, or DMclusters.

### Annotation, enrichment, and network analysis

We annotated cytosines in outlier DMclusters using two reference files (transcriptional and coding-sequence annotations) obtained from the Genes and Gene Predictions Tracks using the UCSC Table Browser for TaeGut2 annotated using the *intersect* function of *bedtools v2.28.0* (Quinlan & Hall 2010). We further assessed if the cytosines annotated as found within transcriptional or coding sequence were functionally enriched for specific gene ontological (GO) categories with the program *g:Profiler* (Raudvere et al. 2019). We included only reference genes with known annotations applied the Benjamini-Hochberg FDR adjustment to the signified threshold (FDR<0.05). We limited the annotation sources for the biological process (BP) GO category, regulatory motifs in DNA as found in miRTarBase, and any information inferred from the human phenotype ontology. Descriptions and child or related GO terms were referenced in *AmiGO 2* version 2.5.12 (last file loaded 2020-03-24). Subcellular localization information on proteins was obtained from the *Compartments* database, and report the locales with the highest confidence (Binder et al. 2014).

Using *StringApp* in *Cytoscape* v3.8.0 (Assenov et al. 2008), we queried human protein databases to generate a network rooted by each gene name with an enriched GO term(s). We used a confidence cutoff of 0.4 and viewed up to 50 interactors in the network. We then retrieved the functional enrichment information for the nodes contained with a network constructed from the gene with an enriched GO term.

### Genetic variant discovery from methylation sequence data

To maximize SNP variation discovery, we merged the paired sequence from hatchlings and fledglings per individual house wren. We then discovered single nucleotide polymorphisms (SNPs) from the merged converted DNA after mapping it to the TaeGut2 reference genome using *BS-SNPer* (Gao et al. 2015), a program that explicitly discovers genetic variation from bisulfite treated sequence data using a dynamic matrix. We used default parameters with respect to frequencies of alleles to confidently identify heterozygous or homozygous sites (minimum of 10% or 85%, respectively), minimum of 15 base quality, minimum and maximum sequence coverage (10-1000), a minimum of two reads per mutation observed, a minimum mutation rate of 0.02 and mapping value of 20. We used the *intersect* function of *bedtools v2.28.0* to determine if SNPs were annotated in any genomic interval of interest.

### Differential methylation analysis and corticosterone data

Following from above, we also used *MACAU*’s binomial mixed model regression analysis to identify differentially methylated sites with corticosterone as the quantitative and continuous predictor phenotype for either the hatchlings with corticosterone measured at day 0 or for fledglings measured at day 15 (Table S1). We included the first two PC coordinates to represent possible batch effects and parental nest identity as covariates, with the relatedness matrix constructed as described above. We followed the annotation, enrichment, and network analysis methods as described above. However, we limited our outliers to cytosines with *p*-values in the lowest 1^st^-percentile of the distribution that also had beta (ß) values in the 1^st^-percentile of either end of the ß distribution. While this threshold is relatively relaxed, we emphasize the enrichment analysis in which significant cytosines established a candidate gene list. A linear regression model (*lm*) was conducted in R (version 3.6) to examine the relationship of outlier cytosine methylation frequencies and corticosterone levels.

### General linear mixed model

We used R (version 3.6) to perform all generalized linear mixed models (GLMMs) with the *lmer* function in the *lme4* package (Bates et al. 2015). We implemented Tukey post-hoc tests using the *emmeans* function in the *emmeans* package (Lenth 2016). All final models met assumptions and significance was defined as α=0.05. We used a GLMM with repeated measures to test if average methylation frequency at day 0 and at day 15 were different within individuals before and after cross-fostering. We included the interaction of treatment (rural to rural, urban to urban, urban to rural, and rural to urban) and time (day 0 or day 15) as fixed effects. Date of capture and brood size were initially included in all models but due to lack of variation and significance, were removed from final models. Individual ID, chromosome number, and nest ID of the genetic and foster parents were included as random effects. We calculated the variance explained by random effects using the package *sjPlot* and model estimates (Lüdecke 2016). We used the Tukey post-hoc multiple comparison tests for the interaction to test whether each treatment group was different from others.

### Ethics statement

Our methods were approved by the UNR’s Institutional Animal Care and Use Committee, and conducted under relevant state and federal permits.

## Results

### Samples and corticosterone summary

We collected 143 blood samples from 86 individual house wrens, with only two samples that failed to provide enough sequence reads for inclusion in this study (sample IDs 07922 and C22.3), resulting in a total of 85 wrens in the study (Table S1). These 85 wrens provided us with 81 hatchlings (R=38, U=43) and 60 fledglings (R=27, U=33) (Fig. 1A). We constructed three datasets to investigate the impact of genetic and environmental contributions to corticosterone levels and methylation levels in: 1) 80 hatchlings from either rural (R) or urban (U) nests; 2) 56 fledglings grouped based on their hatchling (i.e. nest) location of rural or urban to reflect the “genetic” influence on corticosterone and methylation; and 3) the same 56 fledglings grouped based on their fledgling location of rural or urban, regardless of their hatching location, to reflect the influence of their fledgling location on corticosterone and methylation.

Our phenotype analysis here of corticosterone was for a subsample of that previously published (Ouyang et al. 2019). As such, we present briefly a survey of corticosterone to ensure the overall trend was consistent with a smaller sample size (Table S2). Similarly to the previous study, we found no significant difference in corticosterone levels between fledglings at day 15 with respect to their hatching locations (fledglings that hatched from: rural=16.7, urban=17.8, *t*=−0.4, *df*=48.1, *p*=0.6633); rather, their fostered environment post-translocation significantly increased corticosterone in urban fledglings (Table S2). These results support the further investigation of the genetic versus environmental influence on methylation patterns as it pertains to the nest environment and corticosterone levels.

### Methylation summary and outlier sample screen

We obtained an average of 27,891,898 raw reads per individual (total of 141 samples) after filtering for a minimum of 10-fold sequence coverage, with an average of 9,880,751 reads that uniquely mapped to the reference zebra finch genome (Table S1). Post filtering, we obtained 17.9-fold average sequence depth across cytosines (s.d.=7.3x) and all bisulfite conversion rates were high (range=0.981-0.997). We estimated methylation frequency (MF) values for 141 samples at a minimum of 10-fold cytosine sequence coverage for 41,184 cytosines across CG di-nucleotide motifs (n, autosomes=39,692; Z chromosome=1,492). We analyzed three datasets (Fig. 1A) limited to cytosines in the CG motif with a minimum of 10x coverage. The PCA of hatchlings did not reveal clear clustering of samples by hatching environment (Fig. 1B). Methylation data from fledglings, however, revealed a tighter clustering of urban hatched and fledged wrens, spatially distinct from rural hatchlings raised in urban nests (Fig. 1B).

### Urban hatchlings are hypermethylated, with decreased methylation when environments changed

The interaction of time (day 0 or day 15) and treatment on methylation frequency was significant (Table 1). At day 0, rural hatched nestlings had lower average methylation frequencies than urban hatched nestlings (Fig. 2; Tables 1 and 2). Specifically, within-individual changes from day 0 to day 15 showed that average methylation frequencies increased in chicks that stayed in the same parental or foster environment, i.e., moved from rural nests to rural nests (coef=0.05, s.e.=0.001, z=81.3, *p*<0.001) or moved from urban to urban nests (coef=0.1, s.e.=0.001, z=120.4, *p*<0.001). The increase in autosomal methylation frequency from day 0 to day 15 was smaller in RU or absent in UR when hatchlings were moved to foster nests across environments, *i.e*., moved from rural nests to urban nests (coef=−0.005, s.e.=0.001, z=−6.14, *p*<0.001) or moved from urban nests to rural nests (coef=−0.01, s.e.=0.001, z=−22.3, *p*<0.001). Individual ID, chromosome number, and parental and foster nest ID explained very little variation in average methylation frequencies (Table 1).

**Figure 2.**
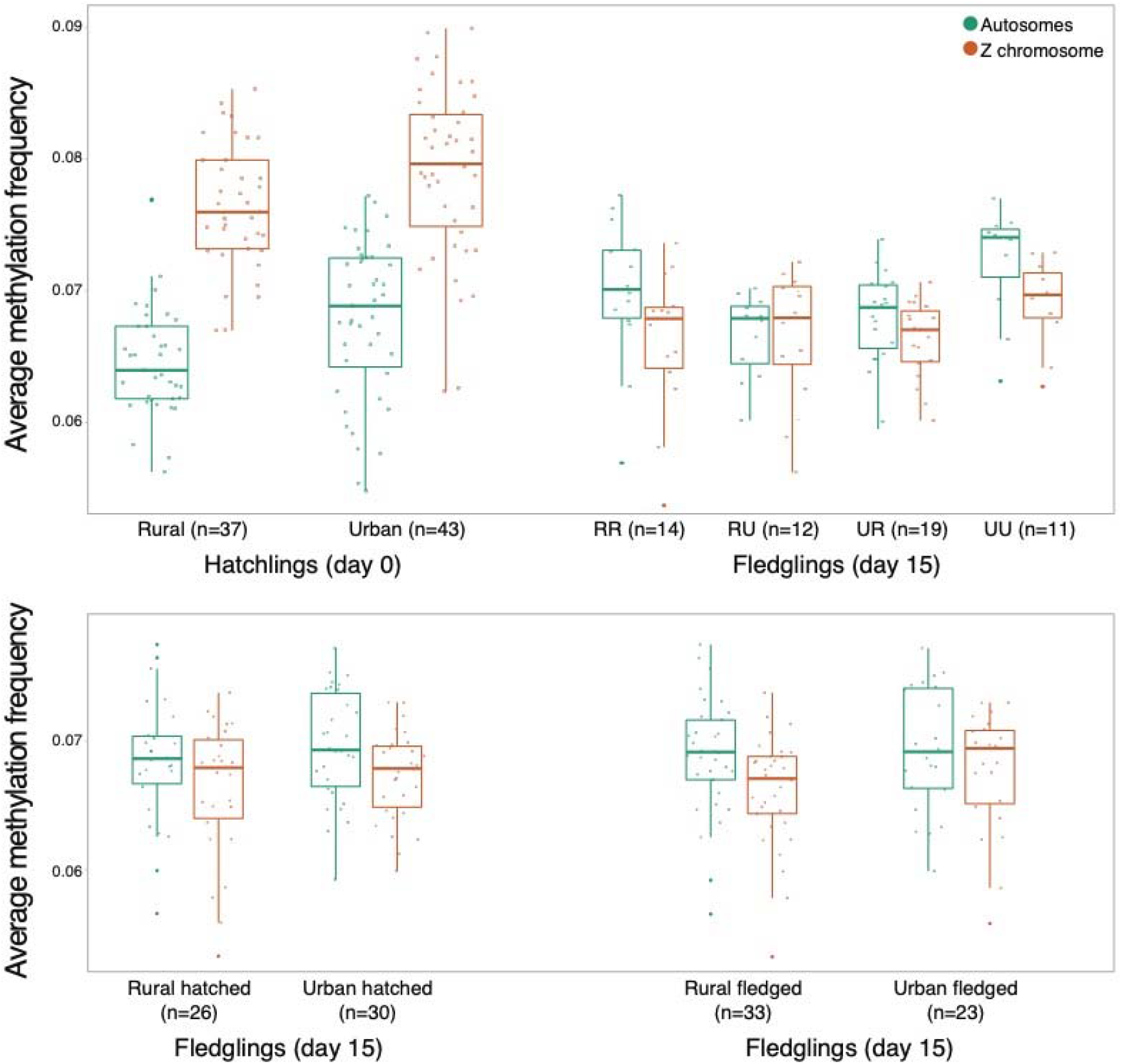
Box-and-jitter plots of methylation frequency of CG-motif cytosines across autosomes and Z chromosomes for hatchlings and fledglings. (Abbreviations: n, sample size; RR, rural hatchling that fledged from rural nest; RU, rural hatchling that fledged from urban nest; UR, urban hatchling that fledged from rural nest; UU, rural hatchling that fledged from urban nest)

**Table 1.** Model estimates for the effect of cross-foster treatment on average methylation frequency of nestling house wrens. Individual estimates are given from summary statistics of the GLMM (binomial for frequency). Random effects include chromosome number, individual ID, and parental and natal nest identity. Time is either day 0 or day 15 for blood sampling. (Abbreviations: R, rural; RR, rural hatchling that fledged from rural nest; RU, rural hatchling that fledged from urban nest; U, urban; UR, urban hatchling that fledged from rural nest; UU, rural hatchling that fledged from urban nest)

| Variable | Estimate | s.e. | z | p-value |
| --- | --- | --- | --- | --- |
| *LMM for corticosterone levels* |  |  |  |  |
| (Intercept) | -2.29 | 0.01 | -183.018 | <0.001 |
| Crossfoster group RU (*reference group RR*) | 0.001 | 0.01 | 0.081 | 0.935 |
| Crossfoster group UR | 0.05 | 0.01 | 3.96 | <0.001 |
| Crossfoster group UU | 0.08 | 0.01 | 5.89 | <0.001 |
| Time (day 15) | -0.05 | 0.00 | -81.32 | <0.001 |
| Crossfoster group RU x Time (day 15) | 0.06 | 0.001 | 56.77 | <0.001 |
| Crossfoster group UR x Time (day 15) | 0.07 | 0.001 | 75.05 | <0.001 |
| Crossfoster group UU x Time (day 15) | -0.04 | 0.001 | 41.12 | <0.001 |
| Random Effects |  |  | variance | SD |
| Chick ID |  |  | <0.001 | 0.089 |
| Chromosome ID |  |  | 0.15 | 0.38 |
| Foster nest ID |  |  | <0.001 | 0.01 |
| Parental nest ID |  |  | <0.001 | <0.001 |

**Table 2.** Estimates from Tukey post-hoc multiple comparisons to test differences among crossfoster groups. Individual estimates are given from post-hoc test on the interaction between time (day 0 or day 15) and cross-foster group. (Abbreviations: R, rural; RR, rural hatchling that fledged from rural nest; RU, rural hatchling that fledged from urban nest; U, urban; UR, urban hatchling that fledged from rural nest; UU, rural hatchling that fledged from urban nest)

| *Post-hoc comparisons of average methylation frequencies at day 0 (hatch date)* |  |  |  |  |
| --- | --- | --- | --- | --- |
| Cross-foster group | Estimate | SE | z | p-value |
| RR-RU | -0.001 | 0.01 | -0.08 | 1.00 |
| RR-UR | -0.05 | 0.01 | -3.96 | 0.002 |
| RR-UU | -0.08 | 0.01 | --5.89 | <0.001 |
| RU-UR | -0.05 | 0.02 | -3.01 | 0.05 |
| RU-UU | -0.08 | 0.02 | -4.19 | <0.001 |
| UR-UU | -0.03 | 0.02 | -1.51 | 0.80 |
| *Post-hoc comparisons of average methylation frequencies at day 15 (~2 days before fledge)* |  |  |  |  |
| RR-RU | -0.06 | 0.01 | -4.56 | <0.001 |
| RR-UR | -0.12 | 0.01 | -9.25 | <0.001 |
| RR-UU | -0.04 | 0.01 | -2.62 | 0.15 |
| RU-UR | -0.06 | 0.02 | -3.54 | 0.01 |
| RU-UU | 0.03 | 0.02 | 1.39 | 0.86 |
| UR-UU | 0.08 | 0.02 | 4.67 | <0.001 |

### Differentially methylated clusters identify environment-associated patterns

With the physical definition that a cluster of differentially methylated cytosines (referred to as dm-clusters) had an inter-cytosine distance ≤40bp, we identified dm-clusters with respect to nest location and age or environment of house wrens (Fig. S1), to determine the relative importance of environment and genetics in shaping methylation patterns across the genome. We outlined the results from each analysis of differential methylation here.

*Behavior and metabolism genes are differentially methylated in hatchlings from different environments.* We identified 7,472 differentially methylated cytosines residing in 2,491 dm-clusters in 80 hatchlings with respect to nests locations in rural or urban landscapes. We found 3,190 cytosines that were hyper-methylated in rural hatchlings (average MF, rural=0.14, urban=0.10), and included cytosines in 42 exons (rural=0.15, urban=0.13), 67 introns (rural=0.16, urban=0.11), 112 promoters (rural=0.13, urban=0.10), and 2,969 cytosines were intergenic (rural=0.14, urban=0.10). We found 4,282 cytosines hyper-methylated in urban hatchlings (average MF, rural=0.17, urban=0.22) across 39 exons (rural=0.12, urban=0.16), 70 introns (rural=0.18, urban=0.23), 143 promoters (rural=0.15, urban=0.19), and 4,030 cytosines were intergenic (rural=0.17, urban=0.22).

We found that 127 dm-clusters contained transcribed or coding sequence of 34 known or 65 suspected gene regions. An analysis for gene enrichment identified 70 ontological terms significantly enriched (FDR adjusted *p*<0.05) for biological process, composed of unique genes populating three broad functional categories: behavior/learning; metabolism and cellular response; and detoxification (Tables 3A and S2A). The most significantly enriched term was mechanosensory behavior (GO:0007638, *p*=0.0028), supported by differential methylation in clusters that contained the genes *CNTNAP2*, *FOXP2*, and *STRBP* (Table 4A). Both *CNTNAP2* and *FOXP2* were associated with hyper-methylated dm-clusters in rural hatchlings (average MF: rural=0.08 and 0.30, urban=0.06 and 0.19, respectively), while the dm-cluster associated with *STRBP* was hyper-methylated in urban hatchlings (rural=0.01, urban=0.04). A network analysis of these three genes revealed two networks with a single edge connection between them. The largest network was centered by *CNTNAP2* and *FOXP2* (40 nodes and 181 edges), with 401 significantly enriched terms that included the nervous system (GO:0007399, FDR *p*=1.95×10^−12^), generation of neurons (GO:0048699, FDR *p*=2.62×10^−12^), and vocalization behavior (GO:0071625, FDR *p*=8.29×10^−12^) (Table S3). The smaller network contained *STRBP* (13 nodes and 28 edges), with a single connection to the *CNTNAP2*/*FOPX2* network (Fig. 3A). This network contained 56 enriched terms, dominated by mRNA binding (GO:0003729, *p*=0.0037), cytoplasm (KW-0963, *p*=0.0041), and RNA processing (GO: 0006396, *p*=0.0043) (Table S4). Finally, we examine if underlying genetic variation could explain differences in methylation within these DMclusters. We found no segregating SNPs across groups of hatchlings (Table S5A).

**Table 3.**
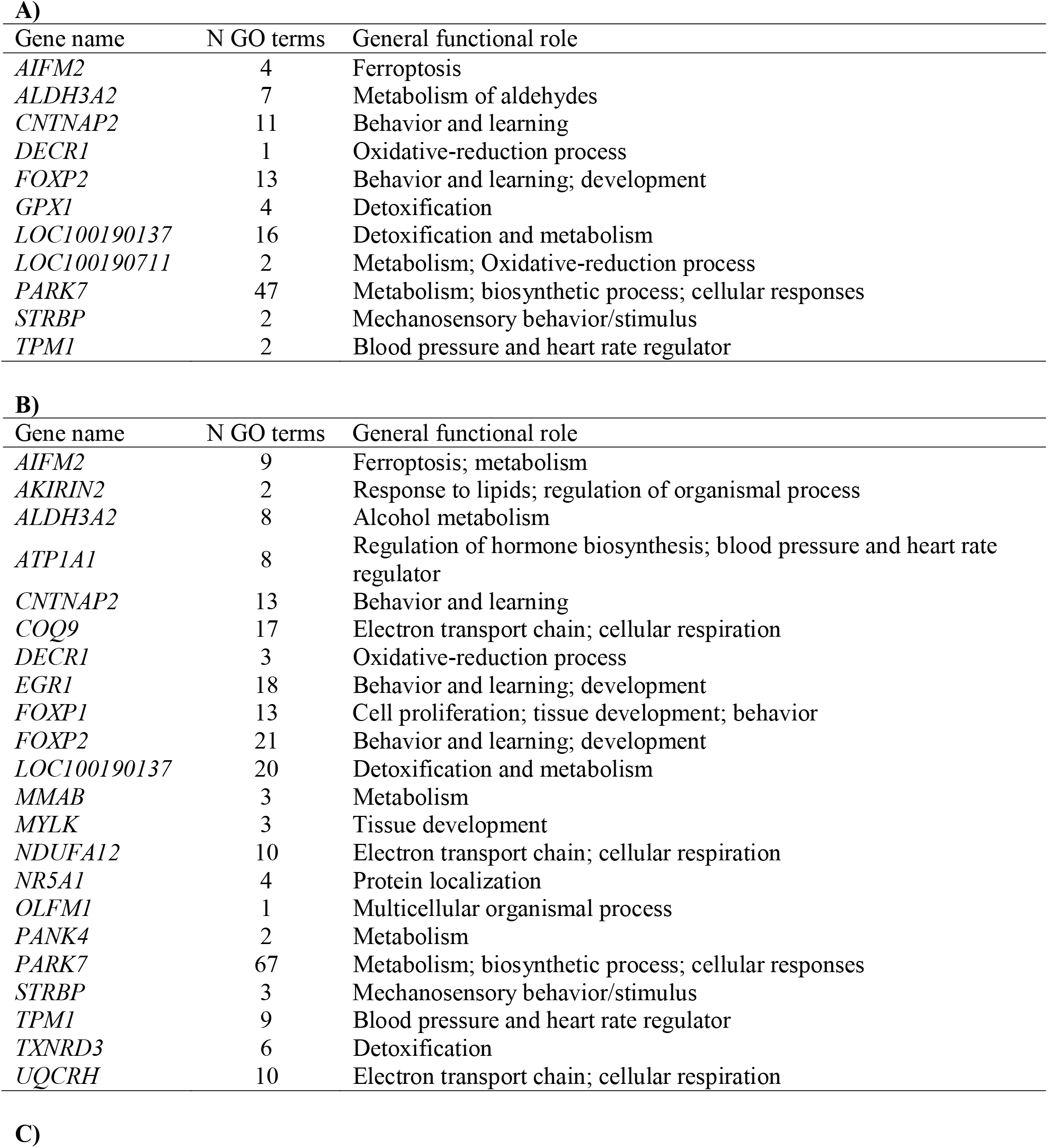
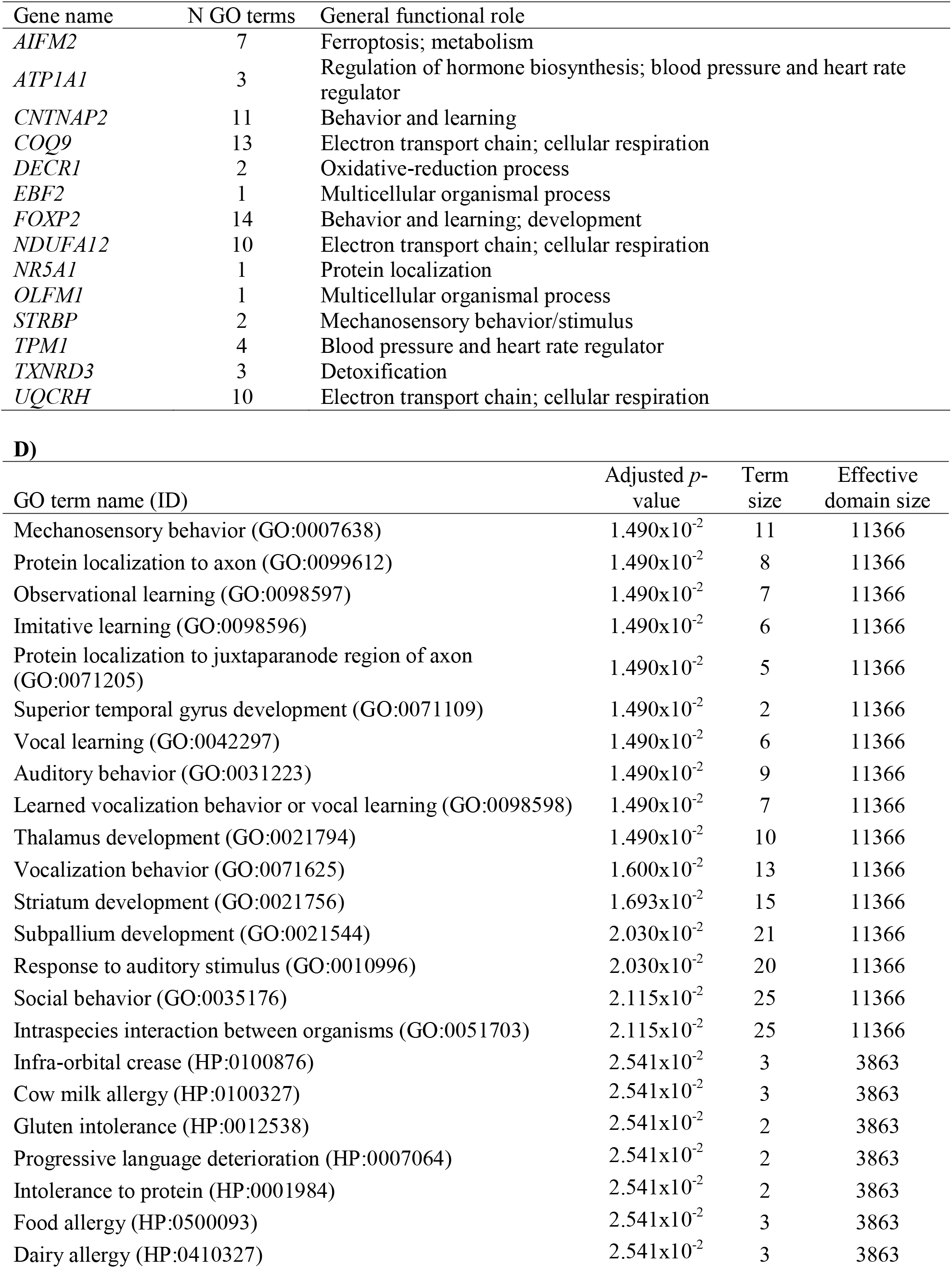
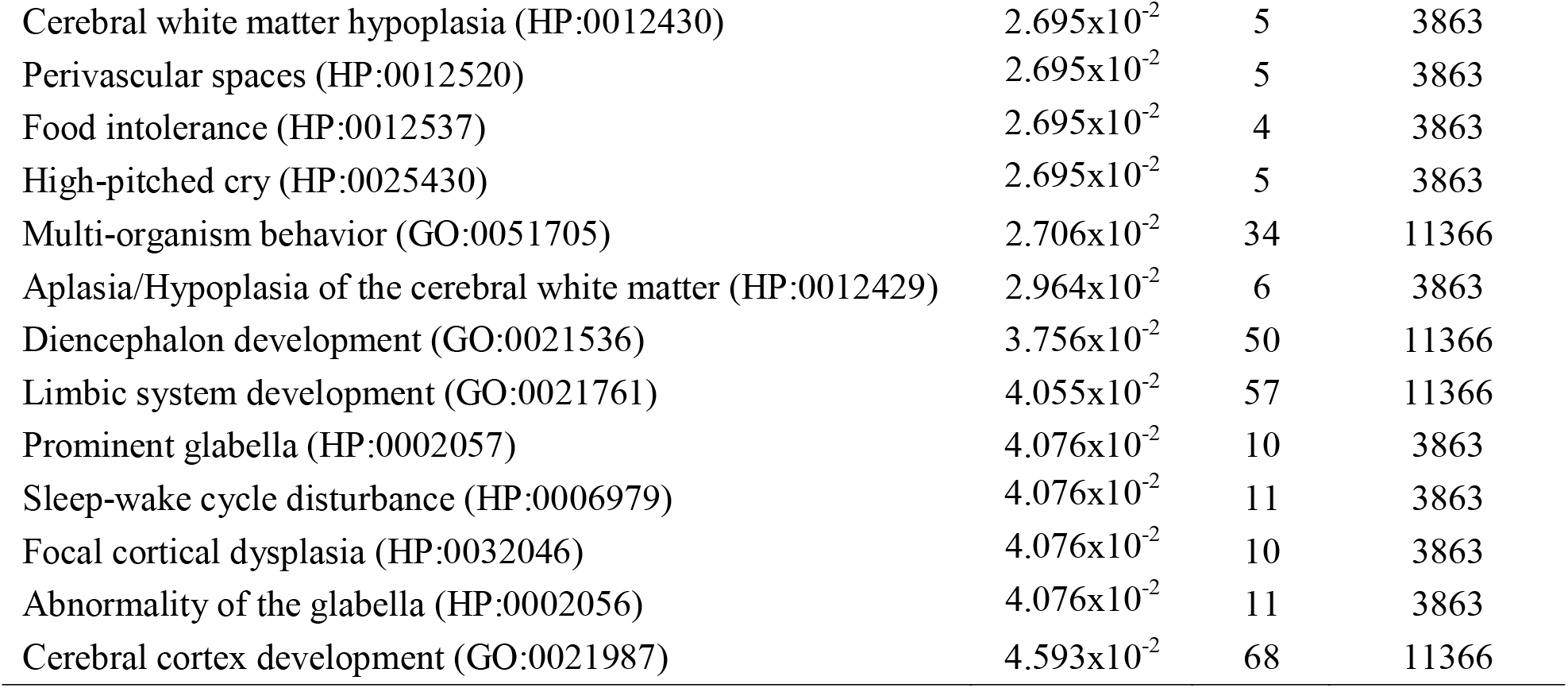
Unique genes identified from the gene ontology (GO) enrichment analysis of three differential methylation data sets with respect to environment (nest location) in **A)** 80 hatchlings (rural versus urban nest locations); **B)** 60 fledglings based on their hatching (not fledgling) nest location; **C)** 60 fledglings based on their fledgling (not hatching) nest location; and **D)** a list of significantly enriched GO terms associated with the cytosines of gene *CNTNAP2* that were differentially methylated with respect to corticosterone measured at day 0 in 80 hatchlings. For details on the GO terms associated with each gene, see Table S2. All GO terms included here were significantly enriched (adjusted *p*<0.05). (Abbreviations: GO, gene ontology; HP, human phenome; N, number)

**Table 4.**
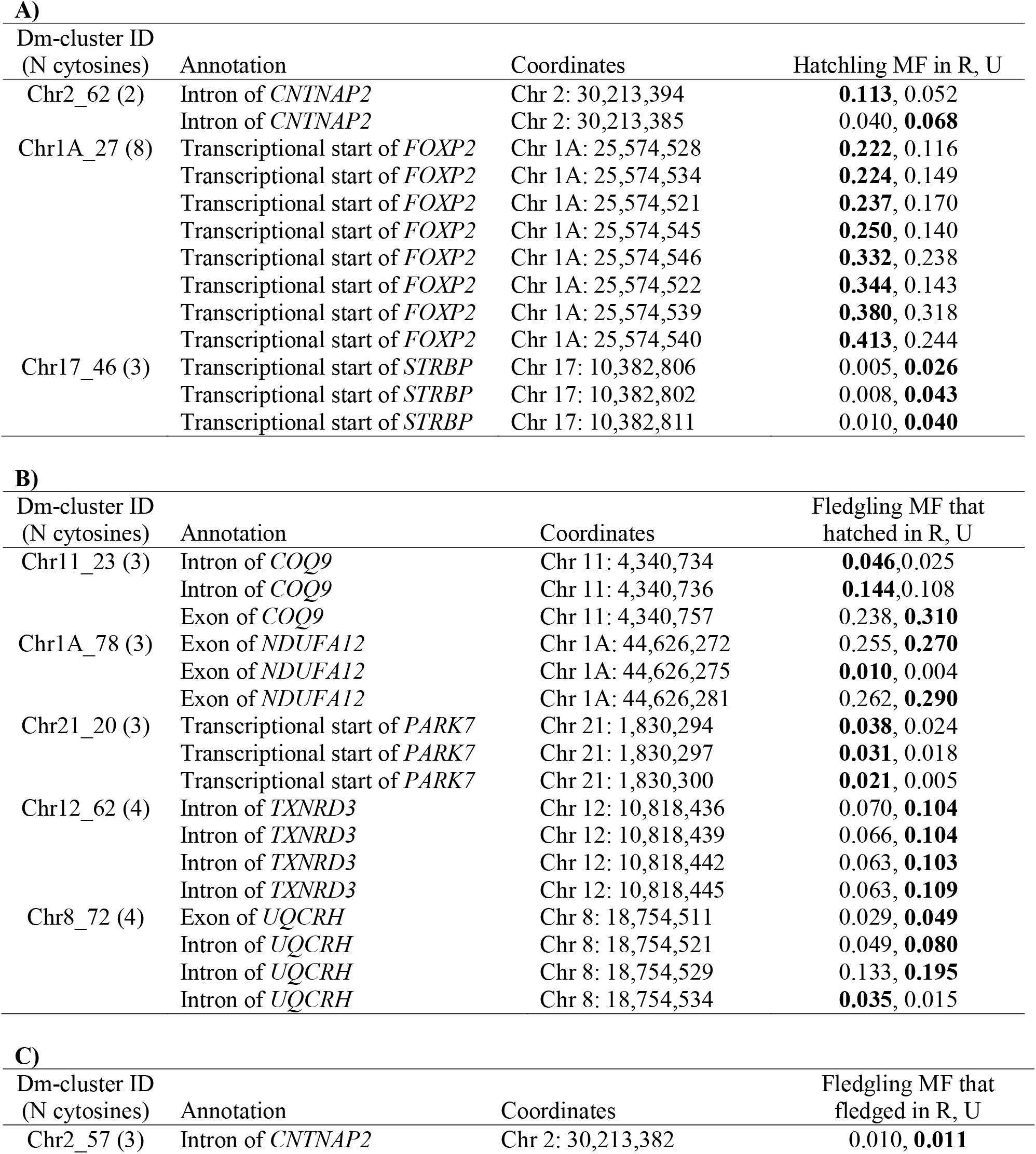
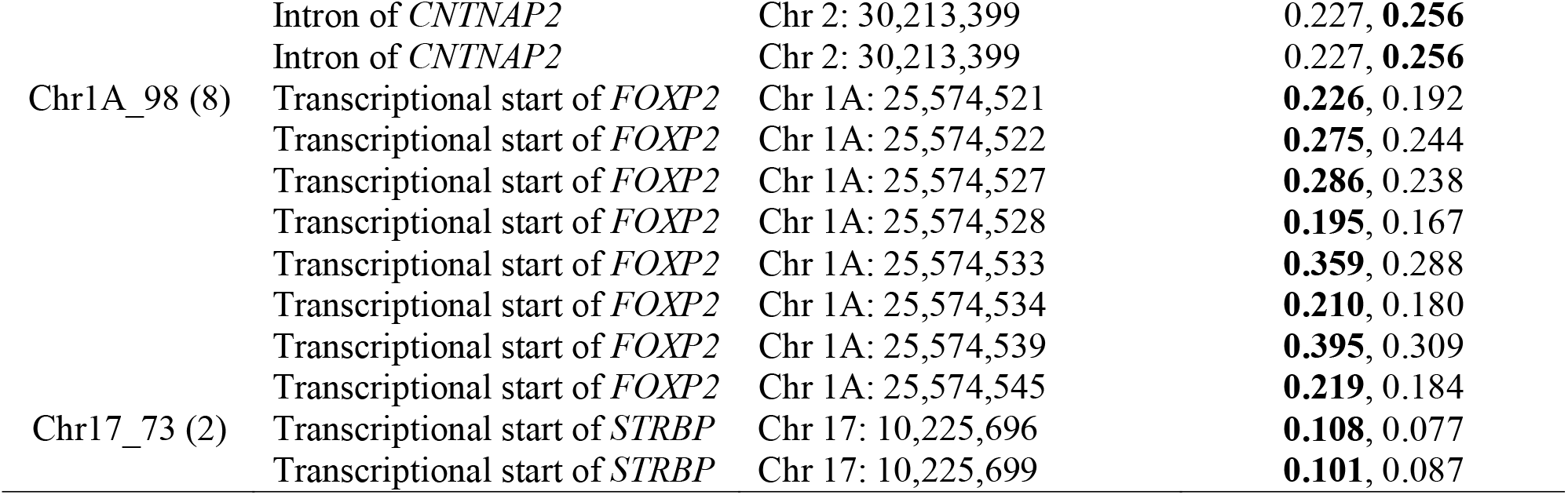
Details for the most significantly enriched Gene Ontology (GO) term for significantly differentially methylated clusters (dm-cluster) with respect to environment (nest location) in **A)** 80 hatchlings (rural versus urban nest locations), **B)** 60 fledglings based on their hatching (not fledgling) nest location, and **C)** 60 fledglings based on their fledgling (not hatching) nest location. All genomic coordinates are provided as chromosome and position mapped to the zebra finch reference genome (TaeGut2). Bolded values indicate the group with hyper-methylation. (Abbreviations: Chr, chromosome; MF, methylation frequency; N, number; R, rural; U, urban)

**Figure 3.**
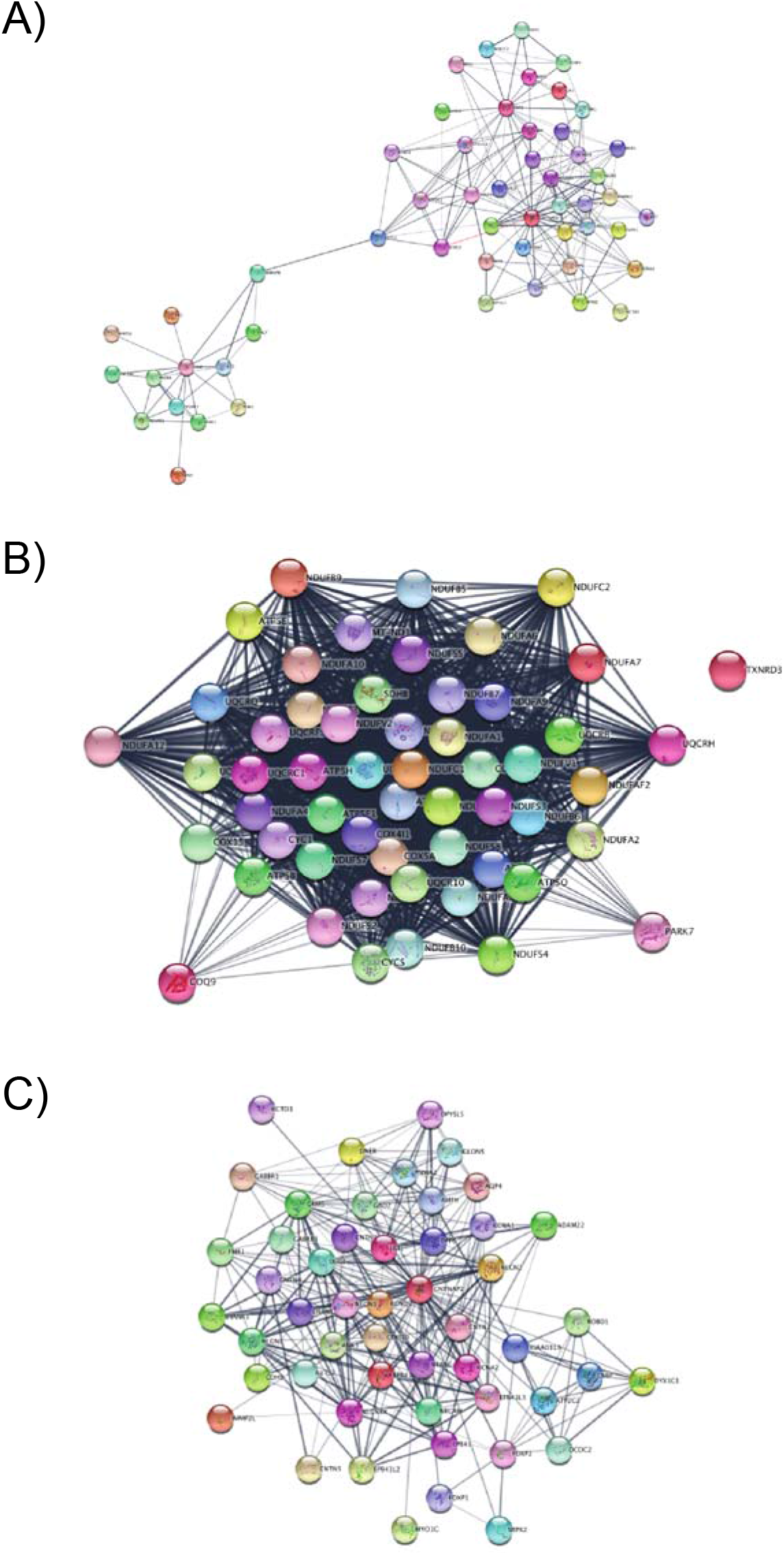
Protein networks for **A)** *STRBP* (lower left) and *CNTNAP2*/*FOXP2* (upper right); **B)** Protein network containing *COQ9, NDUFA12*, *PARK7*, and *UQCRH*, with the single unconnected protein of *TXNRD3*; and **C)** Protein network of *CNTNAP2*, an enriched gene ontology (GO) term with differential methylation associated with corticosterone levels in 80 hatchlings.

Genetic effects on DNA methylation: Cellular respiration gene are differentially methylated in fledglings of different hatching environments. We identified 16,431 cytosines residing in 4,745 dm-clusters with respect to hatchling location of 60 fledglings. We found 7,571 cytosines that were hyper-methylated in fledglings that hatched from rural nests (average MF, rural=0.19, urban=0.14), and included cytosines in 75 exons (rural=0.28, urban=0.24), 129 introns (rural=0.25, urban=0.19), 222 promoters (rural=0.19, urban=0.15), and 7,078 cytosines were intergenic (rural=0.19, urban=0.14). We found 8,860 cytosines hyper-methylated in fledglings that hatched from urban nests (average MF, rural=0.16, urban=0.21) across 117 exons (rural=0.11, urban=0.15), 147 introns (rural=0.19, urban=0.24), 225 promoters (rural=0.14, urban=0.18), and 8,311 cytosines were intergenic (rural=0.16, urban=0.21).

We found that 175 dm-clusters contained transcribed or coding sequence of 45 known or 91 suspected gene regions. An analysis for gene enrichment identified 113 ontological terms significantly enriched (FDR adjusted *p*<0.05) for biological process, made up of cytosines associated with 22 unique genes that populated five broad functional categories: behavior/learning; metabolism and cellular response; cellular respiration; detoxification; and biosynthetic process (Tables 3B and S2B). As found in the hatchlings, the GO term mechanosensory behavior (GO:0007638) was still significant, albeit now ranking as the 5^th^ most significant term (*p*=0.0015). The top four GO terms involved the electron transport chain: mitochondrial ATP synthesis coupled electron transport (GO:0042775, *p*=0.0006); ATP synthesis coupled electron transport (GO:0042773, *p*=0.0006); electron transport chain (GO:0022900, *p*=0.0008); and respiratory electron transport chain (GO:0022904, *p*=0.0010), supported by differential methylation in clusters that contained the genes *COQ9*, *NDUFA12*, *PARK7*, *TXNRD3*, and *UQCRH* (Table 4B). A network analysis of these five genes revealed one large networks containing *COQ9*, *NDUFA12*, *PARK7*, and *UQCRH* (55 nodes and 1,301 edges) while *TXNRD3* was unconnected as a singleton (Fig. 3B). The network contained genes enriched in 151 terms, with the most significant including the respiratory electron transport (HSA-163200, *p*=1.06×10^−101^), Parkinson’s disease (hsa05012, *p*=3.07×10^−99^), oxidative phosphorylation (GO:0006119, *p*=5.03×10^−98^), and the citric acid cycle (HSA-1428517, *p*=1.51×10^−95^) (Tables S6). Additionally, we found no segregating SNPs across groups of fledglings, and thus cannot explain patterns in methylation driven by underlying *cis*-genetic variation (Table S5B).

Environmental effects on DNA methylation: Behavior and metabolism genes are differentially methylated in fledglings from different environments. We identified 9,283 cytosines residing in 2,882 dm-clusters in 60 fledglings assessing environmental influence with respect to nests locations in rural or urban landscapes. We found 4,327 cytosines that were hyper-methylated in rural fledglings regardless of their hatching locations (average MF, rural=0.20, urban=0.17), and included cytosines in 44 exons (rural=0.14, urban=0.11), 71 introns (rural=0.22, urban=0.19), 97 promoters (rural=0.22, urban=0.19), and 4,116 cytosines were intergenic (rural=0.20, urban=0.17). We found 4,956 cytosines hyper-methylated in urban fledglings (average MF, rural=0.23, urban=0.27) across 67 exons (rural=0.20, urban=0.23), 103 introns (rural=0.23, urban=0.27), 141 promoters (rural=0.25, urban=0.28), and 4,645 cytosines were intergenic (rural=0.23, urban=0.27).

We found that 109 dm-clusters contained transcribed or coding sequence of 33 known or 59 suspected gene regions. An analysis for gene enrichment identified 33 ontological terms significantly enriched (FDR adjusted *p*<0.05) for biological process, composed of 14 unique genes across the same five aforementioned broad functional categories: behavior/learning; metabolism and cellular response; cellular respiration; detoxification; and biosynthetic process (Tables 3C and S2C). As found in the hatchlings, the GO term mechanosensory behavior (GO:0007638) was the most significant term (*p*=0.0009), represented by the same three previously described genes *CNTNAP2*, *FOXP2*, and *STRBP* (Table 4C). The difference from hatchling patterns was that rural fledglings were hyper-methylated at *STRBP* (rural=0.10, urban=0.08). Further, *FOXP2* was entirely hyper-methylated in fledglings that fledged from rural nests, regardless of their hatchling location (rural=0.27, urban=0.23). *CNTNAP2* was the only dm-cluster hyper-methylated in urban fledglings (rural=0.15, urban=0.17). The same network interactors and composition was found as in the hatchlings, with a concentration of protein members with functions in behavior and neurodevelopment (Fig. 3C; Tables S3 and S4). We found no segregating SNPs across groups of fledglings, and thus cannot explain patterns in methylation driven by underlying genetic variation (Table S4B).

### Hyper-methylation at a single gene is associated with increased corticosterone levels

A final effort was motivated by the significant corticosterone differences observed between rural and urban environments (Fig. S2). We used the quantitative corticosterone phenotypes to conduct two independent genome-wide association analyses across: 1) 80 hatchlings with methylation data at 57,723 cytosines and corticosterone levels measured at day 0; and 2) 56 fledglings (i.e. not partitioned into subgroups based on cross-fostering) and corticosterone on day 15 for 63,876 cytosines. We found 277 cytosines with significant methylation differences associated with corticosterone levels in 80 hatchlings, with only 10 sites annotated within transcriptional starts or coding sequences (Table S7A). A single gene, *CNTNAP2*, was enriched for functional ontology, with this gene a member of 36 different GO terms (Table 3D). The terms were heavily populated by behavior, brain, and auditory functions (e.g. mechanosensory, observed learning, imitative learning, vocal learning, etc.). A network analysis of *CNTNAP2* revealed one large networks containing 51 nodes and 306 edges (Fig. 3C). The network contained 438 significantly enriched functional terms, with the most significant involved in neurexins/neuroligins (HSA-6794361, *p*=1.17×10^−17^), neuron parts (GO:0097458, *p*=1.86×10^−17^), and neuronal systems (HSA-112316, *p*=5.86×10^−17^) (Tables S8). We used a linear regression and confirmed a significant negative trend of corticosterone (response variable) and methylation at *CNTNAP2* (cytosine at Chr2.30213392; independent variable) across 80 hatchlings (adjusted R^2^=0.03747, *p*=0.04965) (Figs. 4 and S3). We found only two wrens (hatching location, rural=1, urban=1) with SNPs annotated and thus do not explain patterns in methylation at gene *CNTNAP2* (Table S8A).

**Figure 4.**
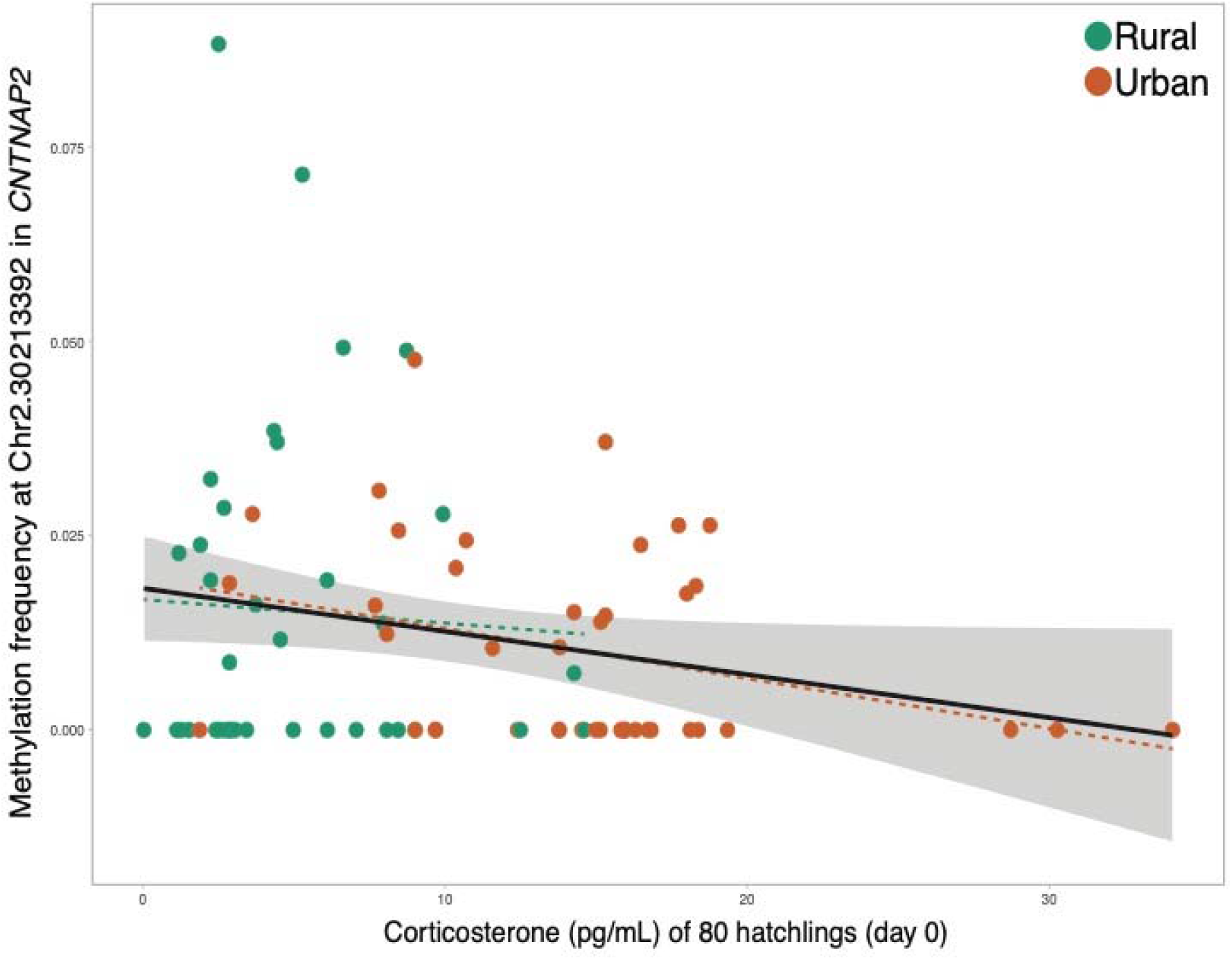
Linear regression of 80 hatchlings and corticosterone levels at day 0 as a function of methylation frequency (MF) at the cytosine (chr2.30213392) in the functionally enriched gene *CNTNAP2*. Trendlines are provided: solid black line is for all hatchlings (corticosterone ~ MF, R^2^=0.05, adjusted R^2^=0.037), whereas dashed lines are for subgroups of hatchlings with respect to their hatching nest environment (rural or urban).

We found significant differences in methylation at 290 cytosines in 56 fledglings, with only 13 sites annotated within transcriptional starts or coding sequences, and none overlapped with the CCA/RDA analysis (Table S7B). Although two cytosines were also round in the CCA/RDA analysis for both the genetic and environmental groupings of the fledgling datasets, none of the 13 outlier genes were functionally enriched.

## Discussion

We used an inter- and intra-environmental crossfoster experiment to analyze the molecular and genetic mechanisms that contribute to early-life phenotypic variation. We found that at hatching, urban offspring had increased average methylation as compared to their rural counterparts. Differential methylation analyses showed that cellular respiration genes were differentially expressed at hatching and behavioral and metabolism genes were differentially expressed at fledgling. Lastly, hyper-methylation of a single gene is associated with increased corticosterone levels. Therefore, both genetic and environmental variance drives DNA methylation patterns.

### Innate DNA methylation differences between urban and rural birds

We found strong genetic signature/architecture with signals of the environment into which an offspring was born. Most of these signals were involved in metabolism, *e.g*., respiratory electron transport, electron transport chain, mitochondrial respirasome, oxidative phosphorylation, and cellular respiration. *EGR1* (also referred to as *ZENK*) is a neural activity-dependent intermediate early gene that has been reported to have decreased expressed in zebra finch that have experienced acute social isolation, whose rescue was mediated through the addition of a partner bird (George et al. 2020). *EGR1* is expressed in the cells that regulate song during singing and in central auditory pathways during hearing, whose expression could result in a long-lasting impact on neuronal cells (Jarvis & Nottebohm 1997; Mello 2002; Reiner et al. 2004; Clayton et al. 2013). Exposure to novel acoustics is known to increase *EGR1* levels in song sparrows and zebra finch, regardless of the social context (Jarvis et al. 1997; Vignal et al. 2005). We also found this gene to be differentially methylation, and part of the GO enriched terms, explaining differences in the genetic influence of fledgling methylation. Urbanization is often associated with increase pollutants, resulting in increased oxidative damage (Isaksson 2010; Salmon et al. 2018) and increased thermal temperatures due to large areas of concrete pavement (Arnfield 2003). Exposure to mercury or methylmercury contaminants resulted in methylation changes at the global and gene level in several species (e.g. bears, mink, rats) (Crews & McLachlan 2006; Bossdorf et al. 2008; Vandegehuchte & Janssen 2011; Basu et al. 2014). Along with differences in nutrient availability where urban areas have less available high-quality food items, it is likely to result in changes in cellular metabolism (Isaksson et al. 2017). There are no studies that we are aware of that have measured endothermic metabolic rates across an urban gradient, but our study points to a promising avenue of hypothesis-drive research, given that some studies have found increased glucocorticoid levels, a hormone that drives glucose metabolism, in cities (Bonier 2012). Although we found strong genetic differentiation between urban and rural birds, we cannot rule out maternal effects, such as incubation behavioral differences or deposition of yolk hormones (Love & Williams 2008; Heppner & Ouyang 2021).

### Effects of urban and rural environments on DNA methylation plasticity

Increased methylation frequencies based on environmental differences have been noted for a number of taxa (Heard et al. 2014; Pedersen et al. 2014; Lea et al. 2016; Sasagawa et al. 2017; Laubach et al. 2018). We also see changes in methylation frequency as young developed (Taff et al. 2019; Venney et al. 2021). In the same environment, offspring increased methylation frequency but when moved to a different environment, offspring decreased methylation frequency. This type of G x E interaction has recently been shown in Chinook salmon in which significant parental and environmental interaction explained phenotypic and methylation frequency variation (Venney et al. 2021).

In fledglings regardless of hatching location, we found strong environmental signatures related to their fledged environment. The most significantly enriched GO subterm was mechanosensory behavior (GO:0007638), which includes auditory behavior (GO:0031223) or a behavioral response to sound. In urban hatchlings, cytosines in the transcriptional element of *STRBP* were hyper-methylated. We hypothesize that urban hatchlings need to have reduced receptivity to an excess of stimulus in the urban environment; thus, increased methylation in the transcriptional start site of this gene could possibly damped the gene’s reactivity. Urban environments are often associated with increased avian song frequency due to increases in anthropogenic noise (Halfwerk et al. 2011). We previously found that our population of urban house wren adults do not respond physiologically when traffic noise was played, which suggests a level of habituation or perhaps reduced sensitivity to sound stimulus in urban environments (Davies 2017). Two additional genes intersected this GO subterm, *CNTNAP2* and *FOXP2*, with hyper-methylation patterns found in rural hatchlings. *CNTNAP2* is a direct *FoxP2* target gene in songbirds, likely affecting synaptic function relevant for song learning and song maintenance (Adam et al. 2017). During periods of enhanced plasticity, such as during development, *FoxP2* influences *CNTNAP2* expression in a linear manner (Adam et al. 2017). Therefore, urban offspring may increase expression of both genes during development as an adaptation to the noisy rearing environment.

### Modification of corticosterone concentrations through DNA methylation of CNTNAP2

As we have found significant and persistent physiological differences in corticosterone expression between urban and rural offspring and adults, we also found a single gene (*CNTNAP2)* that was enriched for functional ontology in urban birds. In other words, hypermethylation of *CNTNAP2* is associated with increased corticosterone levels. GO terms were heavily populated by behavior, brain and auditory functions, *e.g*., observed learning, imitative learning, vocal learning. Urban areas are associated with explorative individuals that exhibit less neophobic responses (Grunst et al. 2019). Stress-reactivity has been shown to be related to explorative behavior and recently has been shown to be related to differentially methylated regions across the genome (Baugh et al. 2012; Taff et al. 2019). To illustrate, females with higher stress resilience, and thus lower feedback in the stress response, had lower overall methylation at a set of differentially methylated regions (Taff et al. 2019). Increased levels of corticosterone have been associated with decreased learning and memory acquisition (Spencer & Verhulst 2007; Monaghan & Spencer 2014). These results combined indicate a relationship between corticosterone levels and learning abilities that may be related to environmental context. Therefore, we hypothesize that decreased *CNTNAP2* expression may result in increased corticosterone levels or vice versa (Ing 2005; Cottrell & Seckl 2009), especially in urban offspring.

We show that DNA methylation exhibits phenotypic plasticity in response to environmental change, such that both parentage and rearing environment influence the methylation status of specific genes. Furthermore, we found strong molecular signatures related to the glucocorticoid phenotype. Our findings are suggestive that DNA methylation can shape the physiological phenotype and is empirical evidence for a mechanism by which individuals thrive in challenging environments. This study provides novel insights for avian epigenetic variation and effects of urbanization, which can generate future empirical studies on specific gene networks and phenotypic traits under selection. Understanding the genetic and environmental basis of local adaptation will be important in predicting species’ responses to urbanization and establishing epigenetic processes as a mechanism for novel environmental acclimation.

## Acknowledgements

JQO is funded by the National Science Foundation (OIA-1738594) and the National Institutes of Health (P20 GM103650). We thank Steve Burgos, Scott Davies, Kelsey Denning, Ryan Fung, Kristiana Hodach, Jeanette Liou, Dante Staten, and David Vijay for valuable field assistance. We thank Caughlin Ranch Housing Association, Reno Park services, and the UNR Mainstation Farm for use of their land.

## Data Accessibility

Sorted BAM files were deposited on NCBI’s Short Read Archive (PRJNA703476).

## Author Contributions

JQO, KVO, KJV, and BMV conceived the project design; BMV and JO collected the data; BMV, RK, and JQO performed the analyses; all coauthors contributed towards manuscript preparation.

