## Supplemental Document for "Reorganization of molecular networks associated with DNA methylation and changes in the rearing environments of the house wren (*Troglodytes aedon*)"

**Supplemental Table S1.** Sample meta-data and sequencing descriptive statistics per individual blood sample (n=144) derived from 86 house wrens. Cross-fostering treatments are represented as the natal nest in which the bird hatches, followed by the next to which it was placed after hatchling and from where it fledged (RR, rural-to-rural; RU, rural-to-urban; UR, urban-to-rural; UU, urban-to-urban). The age of the nest indicates how many days into the nesting season was the nest actively used (1=May 1^st^, 2=May 2^nd^, etc). Corticosterone levels were measured at two time points: hatching at day 0 (Cort0) and fledgling at day 15 (Cort15). Sequence mapping is represented by the: number of raw reads, number of uniquely aligned reads, and percent mappability. Methylation percentage metrics are given for CG or non-CpG cytosines (mCG, mCHG, and mCHH), the average genome-wide bisulfite (BS) conversation rate for phage lambda, the estimates after filtering sites for a minimum of 10x sequence coverage (average coverage and methylation frequency, MF), and the number of SNPs discovered after a 10x sequence filter. (Abbreviations: NA, not applicable)

See file SupplementalTableS1.xlsx

**Supplemental Table S2.** Gene ontology (GO) enrichment results for each of the data sets analyzed for genes with differential methylation with respect to environment (nest location) in **A)** 80 hatchlings (rural versus urban nest locations upon hatching), **B)** 60 fledglings based on their hatching (not fledgling) nest location, and **C)** 60 fledglings based on their fledgling (not hatching) nest location. Effective domain size for all analyses was 11,366.

See file SupplementalTableS2.xlsx

**Supplemental Table S3.** Functional enrichment of the genes included in the network of *CNTNAP2* and *FOXP2*, outlier genes with methylation differentially associated with hatching location (n=80 wrens).

See file SupplementalTableS3.xlsx

**Supplemental Table S4.** Functional enrichment of the genes included in the network of *STRBP*, an outlier gene with methylation differentially associated with hatching location (n=80 wrens).

See file SupplementalTableS4.xlsx

**Supplemental Table S5.** Functional enrichment of the genes included in the network of *COQ9*, *NDUFA12*, *PARK7*, *TXNRD3*, and *UQCRH*, outlier genes with methylation differentially associated with fledgling location (n=56 wrens).

See file SupplementalTableS5.xlsx

**Supplemental Table S6.** Functional enrichment of the genes included in the network of *CNTNAP2*, an outlier gene with methylation differentially associated with corticosterone in 80 hatchlings.

See file SupplementalTableS6.xlsx

**Supplemental Table S7.** Outlier loci identified with an association analysis across **A)** 80 hatchlings with corticosterone levels measured on day 0 and **B)** 56 fledglings with corticosterone levels measured on day 15. Genomic coordinates are provided as mapped to the zebra finch assembly TaeGut2. (Abbreviations: ß, beta value; CDS, coding sequence; TX, transcriptional start)

**A)**

| Cytosine ID (Chr.position) | ß | *p*-value | Annotation | Gene ID |
| --- | --- | --- | --- | --- |
| chr2.30213392 | 6.75x10^-2^ | 1.88x10^-3^ | CDS, TX | *CNTNAP2** |
| chr4A.15540185 | -0.110 | 3.84x10^-4^ | CDS, TX | *LOC100190014* |
| chr18.1012814 | 6.96x10^-2^ | 1.07x10^-3^ | CDS, TX | *LOC100190160* |
| chr20.12440528 | -9.62x10^-2^ | 1.04x10^-5^ | CDS, TX | *LOC100190400* |
| chr5.61597835 | -0.101 | 3.60x10^-4^ | CDS, TX | *LOC100190458* |
| chr24.1567681 | 6.91x10^-2^ | 2.35x10^-4^ | CDS, TX | *LOC100190674* |
| chr19.5243483 | 9.24x10^-2^ | 2.82x10^-5^ | CDS, TX | *LOC100190717* |
| chr3.18752632 | 6.51x10^-2^ | 7.44x10^-4^ | CDS, TX | *LOC100225857* |
| chr5.52214092 | -9.13x10^-2^ | 6.78x10^-5^ | TX | *MIR203* |
| chrZ.43575568 | 9.59x10^-2^ | 1.60x10^-3^ | CDS, TX | *RPL37* |

*This is the only gene with an ontological enrichment.

**B)**

| Cytosine ID (Chr.position) | ß | *p*-value | Annotation | Gene ID | Outlier in CCA/RDA |
| --- | --- | --- | --- | --- | --- |
| Chr3.1494606 | -9.51x10^-2^ | 1.91x10^-3^ | TX | *LOC100190381* |  |
| Chr17.5455051 | 5.72x10^-2^ | 4.38x10^-3^ | TX | *MIR219B* |  |
| Chr26.4902711 | 5.79x10^-2^ | 8.67x10^-5^ | TX, CDS | *FOXP4* |  |
| Chr14.11358858 | -9.45x10^-2^ | 3.84x10^-3^ | TX, CDS | *FSCN2* |  |
| Chr15.2527501 | 6.04x10^-2^ | 3.80x10^-3^ | TX, CDS | *LOC100190104* |  |
| Chr3.19867829 | -6.53x10^-2^ | 4.33x10^-5^ | TX, CDS | *LOC100190175* | Genetic |
| Chr3.19867865 | -8.36x10^-2^ | 3.19x10^-3^ | TX, CDS | *LOC100190175* | Genetic, Environmental |
| Chr2.152589062 | 5.47x10^-2^ | 4.47x10^-3^ | TX, CDS | *LOC100190212* |  |
| Chr5.26594523 | -6.17x10^-2^ | 4.12x10^-3^ | TX, CDS | *LOC100190665* |  |
| Chr4.39701122 | 6.56x10^-2^ | 2.68x10^-3^ | TX, CDS | *MTNR1A* |  |
| Chr2.44385135 | 6.13x10^-2^ | 4.17x10^-3^ | TX, CDS | *NRN1* |  |
| Chr19.7150891 | 8.32x10^-2^ | 2.07x10^-3^ | TX, CDS | *SDF2L1* | Environmental |
| Chr1A.19484843 | 6.18x10^-2^ | 4.18x10^-3^ | TX, CDS | *SELO* |  |

**Supplemental Table S8.** Number of single nucleotide polymorphism (SNPs) annotated with the outlier clusters or genes for **A)** 80 hatchlings and **B)** 56 fledglings. Genomic coordinates are provided as mapped to the zebra finch assembly TaeGut2, represented as chromosome number with the start and stop positions.

See file SupplementalTableS8.xlsx

**Supplemental Figure S1.** The cluster network graph of methylation of an example chromosome Chr1A, for which differentially methylated cytosines (referred to as dm-clusters) were identified between environments and cytosines were proximal (≤40bp apart). Circles indicate hypermethylated and squares indicate hypomethylated differentially methylated sites. (Abbreviations: RR, rural hatchling that fledged from rural nest; RU, rural hatchling that fledged from urban nest; UR, urban hatchling that fledged from rural nest; UU, rural hatchling that fledged from urban nest)

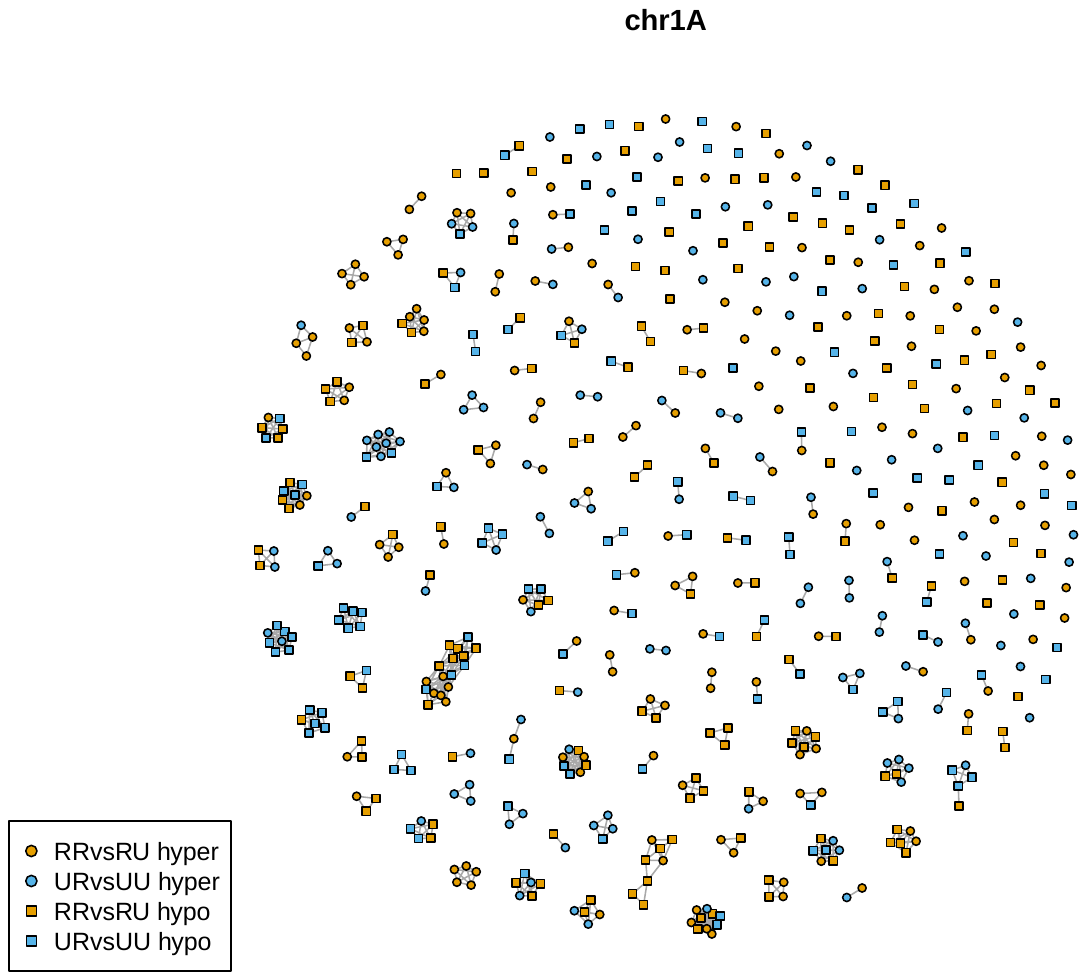

**Supplemental Figure S2.** Boxplots of corticosterone levels (ng/mL) measured in hatchlings (day 0) and fledglings (day 15), as grouped by the crossfostering design. Significant differences between groups were assessed using a Welch Two Sample t-test. Overall corticosterone levels were significantly higher urban hatchlings and fledglings, relative to their rural counterparts (Average corticosterone in ng/mL: hatchlings at day 0 rural=5.0, urban=14.4, Welch Two Sample *t*=-8.2, *df*=68.7, *p*=8.048x10^-12^; fledglings at day 15 rural=12.5, urban=24.2, *t*=-5.2, *df*=46.3, *p*=4.094x10^-6^).

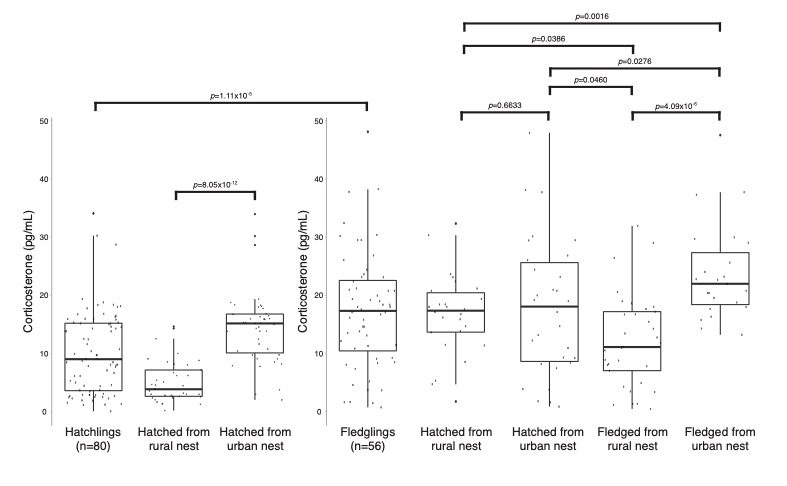

**Supplemental Figure S3.** Residuals of the linear regression model corticosterone (day 0) ~ methylation frequency (MF) at *CNTNAP2* cytosine Chr2.30213392 in 80 hatchling wrens.

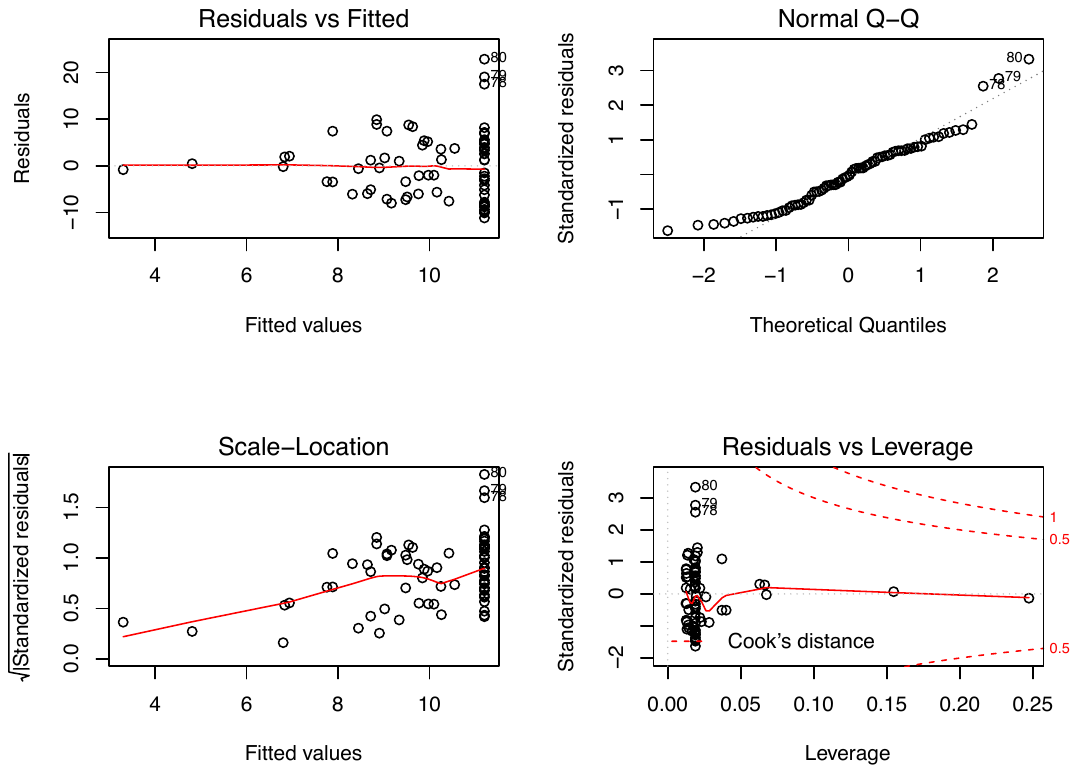
